# TIS Transformer: Remapping the Human Proteome Using Deep Learning

**DOI:** 10.1101/2021.11.18.468957

**Authors:** Jim Clauwaert, Ramneek Gupta, Zahra McVey, Gerben Menschaert

## Abstract

The correct mapping of the proteome is an important step towards advancing our understanding of biological systems and cellular mechanisms. Methods that provide better mappings can fuel important processes such as drug discovery and disease understanding. Currently, true determination of translation initiation sites is primarily achieved by *in vivo* experiments. Here we propose TIS Transformer, a deep learning model for the determination of translation start sites solely utilizing the information embedded in the transcript nucleotide sequence. The method is built upon deep learning techniques first designed for natural language processing. We prove this approach to be best suited for learning the semantics of translation, outperforming previous approaches by a large margin. We demonstrate that limitations in the model performance is primarily due to the presence of low quality annotations against which the model is evaluated against. Advantages of the method are its ability to detect key features of the translation process and multiple coding sequences on a transcript. These include micropeptides encoded by short Open Reading Frames, either alongside a canonical coding sequence or within long non-coding RNAs. To demonstrate the use of our methods, we applied TIS Transformer to remap the full human proteome.

## 1 Introduction

Translation is the synthesis of proteins from messenger RNA (mRNA). The nucleotide sequence of mRNA not only encodes proteins through its codon structure, but also influences other factors, such as the location and efficiency of the translation process (Wilkie et al. 2003). The complexity of the nucleotide sequence is highlighted by the variety of gene products, playing roles in multiple molecular processes. Partly due to this complexity, existing mappings of the genome, transcriptome, and proteome are still mostly based on the combination of experimental data with statistical tests. Genome annotation platforms such as Ensembl, NCBI and USCS rely largely on sequence alignment methods (Aken et al. 2016; Thibaud-Nissen et al. 2016). Flaws of such systems are inherent to the knowledge at hand, where biases and errors are known to be propagated with time (Fields et al. 2015). Subsequently, changes in existing annotations are made as new data on gene products or molecular mechanisms become available.

A comprehensive understanding of translation in combination with the design of high performing machine learning approaches can advance the identification of novel proteins. In recent years, machine learning models have gained attention due to their ability to attain high performances when sufficient data is available. Prior work has attempted to predict TISs sites solely based on intronless transcript sequences (Zien et al. 2000; Chen et al. 2014; Kabir et al. 2015; Zhang et al. 2017; Zuallaert et al. 2018; Kalkatawi et al. 2019; Goel et al. 2020). Deep learning techniques have also gained success due to their automated feature learning (Eraslan et al. 2019), as can be seen through the application of neural networks on sequence and omics data. Zhang et al. (2017) use a combination of convolutional and recurrent layers to process a transcript sequence of fixed length around the TIS. Similarly, Zuallaert et al. (2018) and Kalkatawi et al. (2019) have applied convolutional neural networks to determine the location of TISs. Although the predictive performance of these models still poses restrictions on their wider applications, value from these studies is derived from insights gained on the underlying decision-making process.

Deep learning techniques have the unique strength of uncovering complex relationships in big data. Standard machine learning methods and traditional neural network architectures map relations of the input features to the target label. These relations are, for most architectures (e.g. linear regression, fully-connected layers, convolutions), explicitly mapped as trained weights that determine how information is combined in each layer. In contrast, attention methods determine the relevancy between two pieces of information based on their information embedding and (relative) positioning. Weights applied to recombine information at each layer are calculated on the fly for each new set of inputs. For many natural language processing tasks, this approach has proven superior (Cheng et al. 2016; Parikh et al. 2016; Vaswani et al. 2017). Similar to natural languages, information within biological sequences is complex and convoluted. Some recent studies have successfully shown the efficacy of transformer networks on genome data, such as proposed by Zaheer et al. (2020), for the prediction of promoter regions and chromatin profiling. Ji et al. (n.d.) introduce DNABERT, a transformer model pre-trained on genome data that can be fine-tuned for a specific tasks such as the annotation of promoters or splice sites.

The computational cost of attention imposes limitations on the maximum input length of sequential data. This cost scales quadratically with respect to the length over which attention is calculated. This has traditionally limited the maximum sequence length of previous studies at 512 units (Ji et al. n.d.; Vaswani et al. 2017). However, several recent studies have focused on overcoming this limitation through the use of mathematical approximation techniques that save on computational cost, allowing the attention mechanism to be applied over larger sequences (Wang et al. 2020; Xiong et al. 2021; Choromanski et al. 2021).

In this work, we propose TIS Transformer (see Figure 1), a tool for the identification of TISs using the full processed transcript sequence as input. Our technique uses one of the proposed scaling solutions for computing attention (Choromanski et al. 2021), where transcripts up to a length of 30,000 nucleotides can be processed at single nucleotide resolution. Models are trained and evaluated on the processed human transcriptome, excluding introns and intergenic regions. First, we benchmark our method against previous studies, where we show state-of-the-art performances. Afterwards, we provide an in-depth analysis on the current state of applying learning methods for the annotation of TISs on the full human transcriptome. We show our approach to achieve notable performances in general, where learning and evaluation approaches are mostly hindered by low quality annotation. We believe this approach to be of substantial benefit to the community, where it can aid the discovery and annotation of novel proteins, thus building state-specific proteome maps, or the engineering of transcripts.

**Figure 1:**
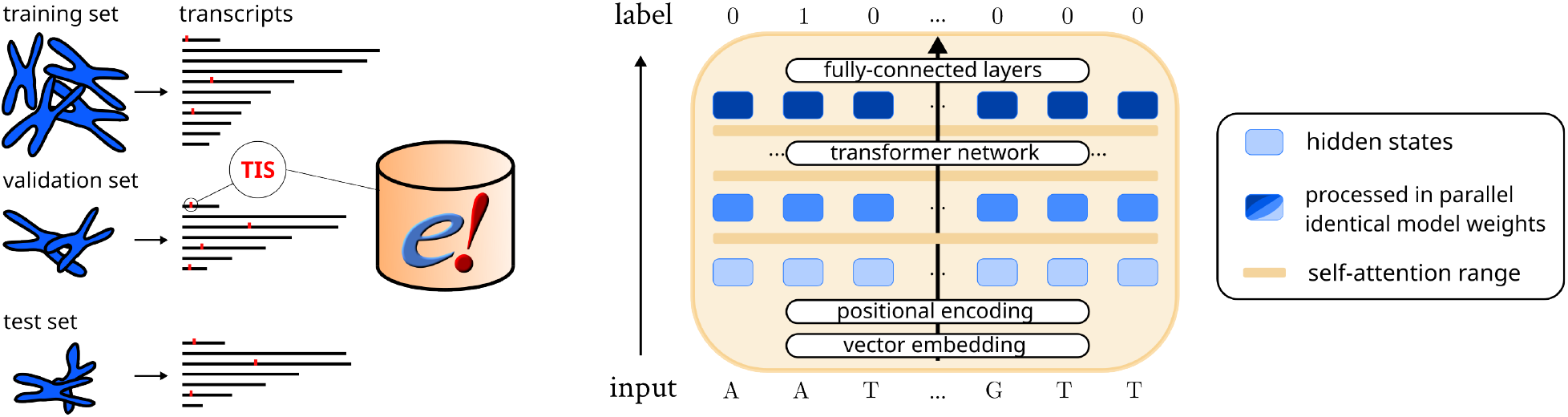
Schematic of the data and model set-up. (**left**) The Ensembl annotation (version 107) is used to determine transcript sequences and translation initiation sites (TISs). Transcripts are grouped by chromosome to create a training, validation and test set. (**right**) The performer model allows processing of full transcript sequences, evaluating data through the layers in parallel to obtain model outputs at each position. The model architecture can handle varying input lengths, as identical model weights are applied to transform the data. Through self-attention, sequential information from any site on the transcript can be queried by the model to determine the presence of TISs at any position.

## 2 Material and Methods

### 2.1 Training objective

The goal of the study is to create a predictive model that is able to detect the occurrence of coding sequences (CDS) within the transcriptome. Rather than predicting a paired TIS and translation termination site (TTS), we decided to achieve this goal through the detection of TISs, as it poses a more simple optimization and decision making problem. TTSs have been determined post-hoc through detection of the first in-frame stop codon. As such, the model is optimized to perform binary classification between TISs and non-TISs for each position of the human transcriptome. We provide all code material and results as part of the study.

### 2.2 Model input and output

The model processes the full sequence of the transcript to impute the presence of TISs at each position (see Figure 1). The input of the model is represented by 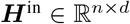, where *n* < 30,000 and *d* the dimension of the hidden states. 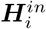 denotes the nucleotide embedding (A, T, C, G, N, [START], [STOP]) at position *i* of the transcript (thymine is used to proxy uracil throughout the paper). The input embeddings are trained as part of the optimization process. The [START] and [STOP] tokens are used at before the beginning and after the end of the transcript, relatively. Identical in dimensions to the input embeddings, the output 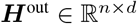 of the transformer network is processed by a pair of feed-forward layers which result in a model output at each nucleotide position of the transcript. The loss is computed from predictions at inputs ∈ {*A, T, C, G, N*}.

### 2.3 Transformer architecture

The transformer architecture is a recent methodology that implements (self-)attention for a group of inputs. Through positional encodings, a sequential ordering can be imposed between these inputs. The attention mechanism is is applied to determine the flow of information (i.e. applied weights to combine information) based on the information itself. The transformer network is characterized by several sequential layers with identical design.

The principal element is the use of self-attention, performed by a module called the attention head. Here, a query, key and value matrix are derived from the input *X* in each layer:

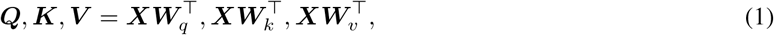

where ***Q, K***, 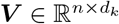. For the first layer, ***X*** equals ***H***^in^. Multiplication and subsequent normalization between ***Q*** and ***K***^τ^ returns an *n* × *n* matrix, whose values determine the flow of information to (e.g. (***QK***^τ^)_*i*,:_) and from (e.g. (***QK***^τ^)_:,*i*_) each hidden state through multiplication with ***V***. For each attention head, a matrix ***Z*** is calculated, holding the combined information for each input hidden state.

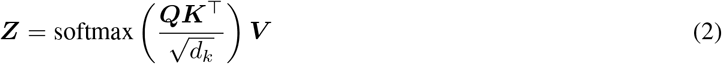

where 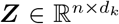. Multiple sets of ***W**_q_, **W**_k_, **W**_v_* make it possible to route multiple types of information from **X** in each layer. The outputs ***Z*** that are derived from different attention heads are concatenated and transformed to fit the dimension size of the input ***X***, with which they are summed (i.e. residual connection). Each layer is built using the same components, albeit unique weights. The architecture allows processing of transcripts of different lengths, as the same weights are applied to calculate the ***q, k*** and ***v*** vectors for each input hidden state (independent of position). These are calculated in parallel for each layer and represented as rows in the matrices ***Q***, ***K*** and ***V***.

To allow long-range attention over full transcripts, we used a recent innovation in calculating attention introduced by Choromanski et al. (2021). Equation 2 is exchanged with the Fast Attention Via Positive Orthogonal Random Features (FAVOR+) algorithm, which utilizes random feature maps decompositions for approximation in order to obtain a computational complexity that scales linearly with respect to the length of the sequence.

### 2.4 Data sets and model optimization

No single data set has previously been designed to function as a benchmark standard when evaluating the prediction of TIS sites. The Ensembl assembly of the Human genome (GRCh38.p13; release 107) was selected to provide the transcript sequences and label the known TISs. The full human genome comprises 96,655 (38.49%) protein-coding and 154,466 (61.51%) non-coding transcripts. These constitute a total of 431,011,438 RNA nucleotide positions, 96,655 of which are positively labeled as a TIS (0.022%), indicating an extremely imbalanced data set. The training, validation, and test set are allocated in accordance to chromosomes, thereby grouping transcript isoforms and proteoforms together. Transcripts longer than 30,000 nucleotides (17 instances) are excluded due to memory requirements. When remapping the full human proteome, six models were trained to sequentially use different sets of chromosomes as test, validation, and train data (Supplementary Table A2 and Figure 1). The annotations on the test sets, constituting one sixth of the chromosomes for each model, are used to obtain a complete re-annotation. To perform the benchmark, several additional constraints were required to ensure all methods were trained and evaluated utilizing the same data (see Supplementary Files). To illustrate, these exclude non-ATG positions and positions at the edge of transcripts that cannot be parsed by one or more of the evaluated methods. The benchmark was performed using chromosome 1, 7, 13, 19 as testing data and chromosome 2 and 14 as validation data. This results in train, validation and test sets of 3,608,307, 641,264 and 1,069,321 candidate TIS positions, respectively. For all instances, training was stopped when a minimum loss on the validation set is obtained. All reported performance metrics are obtained on the test sets. Before benchmarking our method with other studies, hyperparameter tuning was performed (see Supplementary Files for more details on this process). In general, it was observed that the performance of the model was not correlated substantially with any single parameters, but with the total number of model parameters. The number of model parameters is influenced by the dimension of the hidden states (***H***), number of layers, dimension of the attention vectors *q, kandv* and number of attention heads. Through hyperparameter tuning, two model architectures were selected to benchmark against previous approaches, TIS Transformer and TIS Transformer L(arge), each featuring 118K and 356K model parameters. Supplementary Tables A4–5 list the details of each model architecture.

## 3 Results

The study is separated in two main parts. First, we perform a benchmark to compare our method with previous studies, and find our method to achieve state-the-art performances. Afterwards, we remap the human proteome using our approach for a more in-depth analysis on the ability of our models.

### 3.1 Benchmark

A multitude of studies exist that apply machine learning techniques for the prediction of TISs (Supplementary Table A1) (Zien et al. 2000; Saeys et al. 2007; Chen et al. 2014; Kabir et al. 2015; Goel et al. 2020; Zhang et al. 2017; Zuallaert et al. 2018; Kalkatawi et al. 2019; Wei et al. 2021). Two main approaches exist that model sequence information to the occurrence of TIS sites: support vector machines and neural networks. Problematically, the computational cost for support vector machines scales quadratic with the number of samples evaluated. As such, given tools are impossible to be applied on the full genome. To illustrate, the maximum data set size applied by previous studies is 13,503 samples, less than 0.01% of the positions on the full transcriptome. The limited data size allowed to train these models is an important disadvantage, where previous studies show that support vector machine implementations are consistently outperformed by neural network approaches (Zuallaert et al. 2018; Wei et al. 2021; Kalkatawi et al. 2019).

Candidate TIS positions are identical for the training and evaluation of different approaches to ensure a fair comparison. The construction of the training, validation and test set is discussed in more detail in Section 2.4 of the Supplementary Files. We rely on previous results that show listed neural networks approaches to outperform support vector machines, as proposed data set sizes cannot be applied to train support vector machines. Furthermore, we were unable to incorporate the method DeepTIS, as proposed by (Wei et al. 2021), due to missing details on the model architecture in the published paper. See the Supplementary Files for more information on, and an extended discussion of, the individual methods and the benchmark.

Results show the TIS Transforer to largely outperform previous methods (Table 1). As TIS Transformer L(arge) performs better on the benchmark data set, it is the model architecture has been applied to remap the human proteome.

**Table 1:** Benchmarked performances on a set of available tools. For each study are given: the year of publication, the nucleotide input size parsed by the machine learning model, the total number of trainable parameters, the training time until convergence, and the ROC and PR AUC scores. Note that methods applying support vector machines could not be evaluated due to the size of the data, nor DeepTIS due to missing information. The total number of transcript positions used to train, validate and test each method count to a total of 3,608,307, 641,264 and 1,069,321 positions.

| Model name | Year | Input size | parameters | Train time | ROC AUC | PR AUC |
| --- | --- | --- | --- | --- | --- | --- |
| DeepGSR (Kalkatawi et al. 2019) | 2019 | 600nt | ~181M | ~100h | 0.9485 | 0.2110 |
| TITER (Zhang et al. 2017) | 2017 | 203nt | ~431K | ~24h | 0.9617 | 0.3770 |
| TISRover (Zuallaert et al. 2018) | 2018 | 203nt | ~240K | ~3h45 | 0.9650 | 0.4495 |
| TIS Transformer | 2022 | Transcript | ~118K | ~10h | 0.9970 | 0.7628 |
| TIS Transformer L | 2022 | Transcript | ~356K | ~6h | 0.9971 | 0.7684 |

**DeepTIS** (Wei et al. 2021): See supplementary files on missing details on its architecture.
**SVM** (Zien et al. 2000; Chen et al. 2014; Kabir et al. 2015; Goel et al. 2020): Training of support vector machines is not tractable on large data sets.

### 3.2 Evaluation of the remapped proteome

#### 3.2.1 Performance evaluation

To perform an in-depth analysis on the model predictions of the full human transcriptome, a cross-validation scheme is used (see Section 2.4 Supplementary Table A2). The averaged area under the precision-recall curve (PR AUC) score of the six models trained to remap the proteome is 0.839. These scores underline the capability of the TIS Transformer to predict with a high recall and precision given the extremely imbalanced setting, in turn reflecting the usefulness of the model to be applied to predict sites of interest on the full transcriptome. The output probabilities of the negative and positive samples can be clearly distinguished as they are concentrated at 0 and 1, respectively (see Figure 2a). Similarly, this can be observed when evaluating the model outputs at the single transcript level (e.g. see Figure 4).

**Figure 2:**
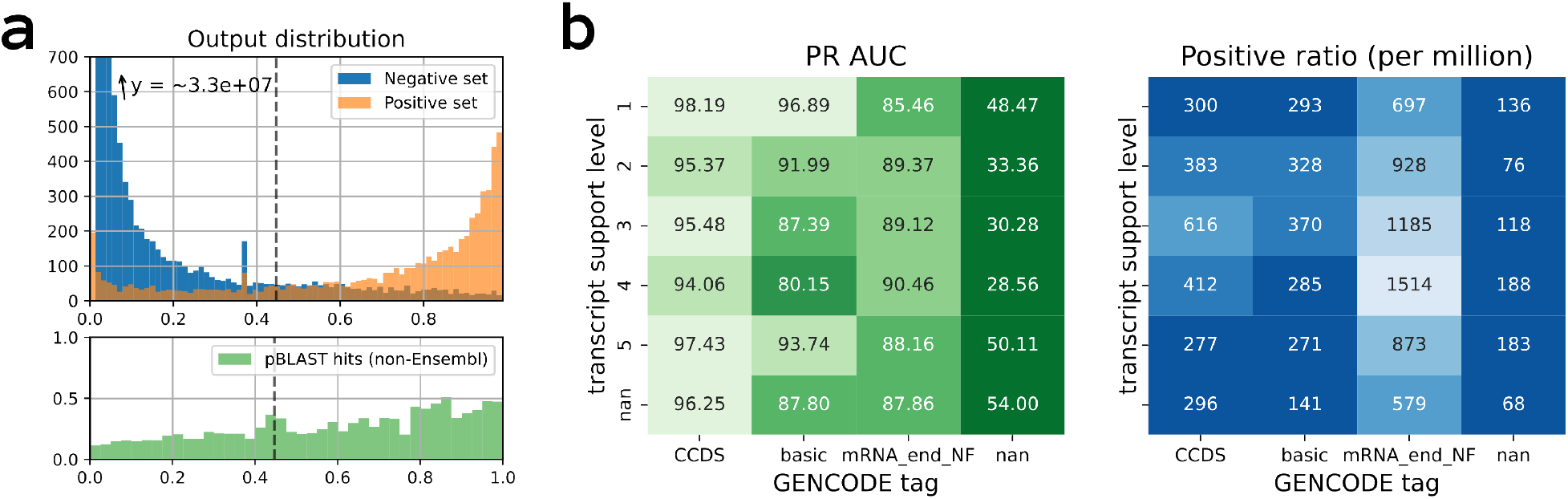
Model performances on predicting translation initiation sites. (**a**) (top) A histogram of the model output predictions when evaluated on chromosome 2 (test set). Transcript sites are divided in a negative (blue) and positive (orange) set according to the annotations provided by Ensembl. The dotted line represents the threshold that an equal number of positive model predictions as provided by Ensembl. (bottom) The resulting coding sequences of predicted TISs were evaluated against UniProt using pBLAST. Shown are the fraction of coding sequences returning a good match in relation to the model output score. Only TISs not annotated by Ensembl (i.e. negative set) were considered. (**b**) Model performances were binned according to transcripts properties. PR AUC performances (left) and the ratio of positive samples in each group (right) are obtained by binning the transcripts according to transcript support level and other properties given to the annotated translation initiation site or transcript (if any, otherwise nan).

**Figure 3:**
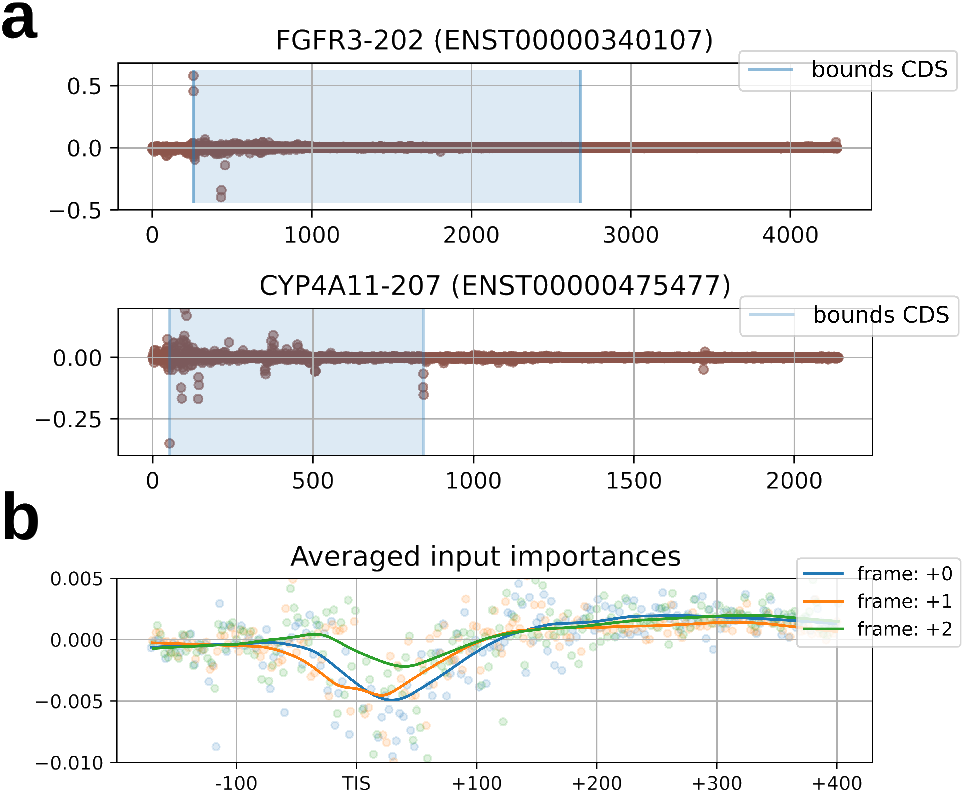
Attribution scores of input nucleotides which reflect the importance of each nucleotide towards predicting the annotated TIS site. (**a**) Scores shown are given by the model for the translation initiation site on position 257 of FGFR3 (ENST00000340107) and at position 52 of CYP4A11 (ENST00000475477). (**b**) Attribution scores for the positions surrounding the translation initiation site (TIS) averaged for all top ranked predictions. a rolling average is given for the three reading frames.

**Figure 4:**
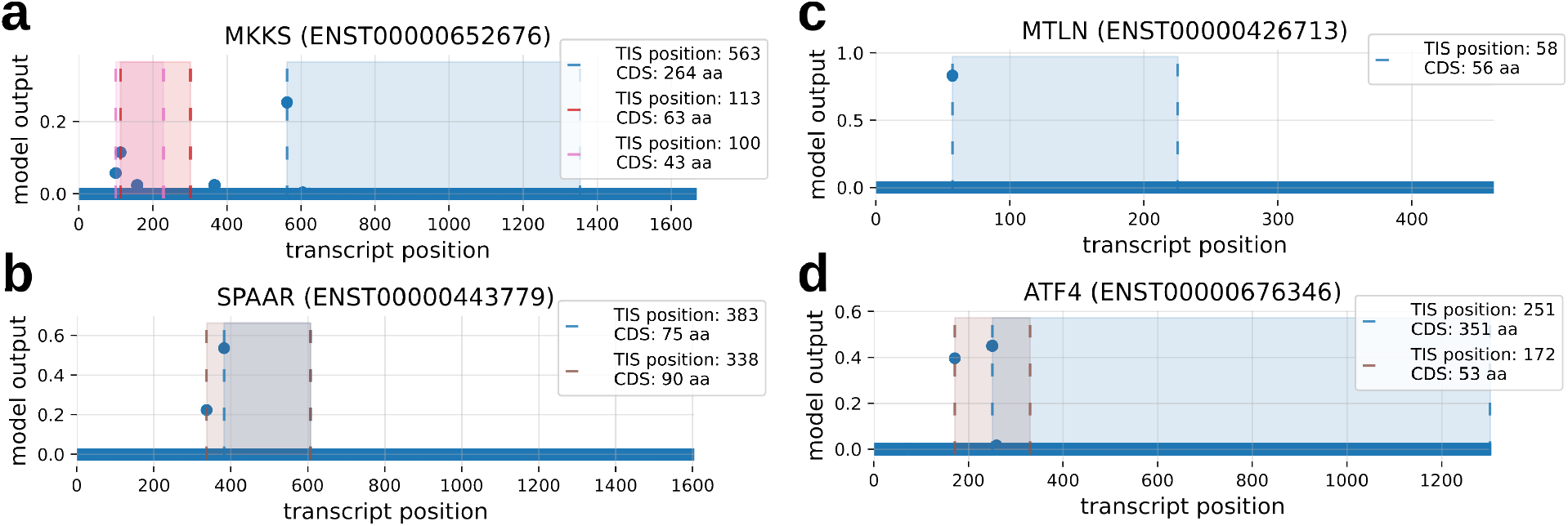
Example transcripts with predicted micropeptide expression. Shown are the model outputs (y-axis) for each position of the transcript (x-axis). For high TIS predictions, the bounds (striped line) and area of the resulting CDSs are shown. Given are the predictions for the transcripts (**a**) MKKS (ENST00000652676), (**b**) SPAAR (ENST00000443779), (**c**) MTLN (ENST00000426713), and (**d**) ATF4 (ENST00000676346).

Several patterns exist between the model performance and data properties as obtained by GENCODE (Figure 2b). Some properties are linked to the transcript, such as the transcript support level, which quantifies the support for a transcript to exist based on available *in vivo* evidence (Yates et al. 2016). Other discussed properties relate to whether these TISs are part of the consensus coding sequence database (CCDS), whether the transcript and resulting coding sequence is identified as a representative sample for the gene (basic), and whether the mRNA end could be confirmed or not (mRNA_end_NF). Supplementary Figure A7 includes more information for each group including the ROC AUC scores and number of nucleotide positions (samples) in each group.

We observe several patterns of special interest. First, the performance of the model is correlated to the transcript support level for the bulk of annotations (CCDS, basic). Only the group with the lowest transcript support level (i.e. 5) shows better performances than expected based on its ranking. As this group of transcripts is described as ’no single transcript supports the model structure’, we hypothesise their presence to be supported by other factors such as computational pipelines, which can be expected to have good performance. Second, the model performs worse for the group of transcripts where no tags are given to the transcript or TISs. These tags include the validation of TISs by other parties such as the CCDS initiative. Third, the differences in performance between groups is largely uncorrelated to the ratio of positive samples along rows and columns, which could otherwise have been the driving factor for differences in PR AUC scores.

Figure 2a shows the relation between the model output and the percentage of resulting CDSs having been observed by previous studies. These show a clear correlation between the model certainty and the percentage of positive matches. These results were obtained by querying CDSs resulting from high-ranking predictions that were not previously annotated by Ensembl against Swiss-Prot and TrEMBL (mammals) using pBLAST. Section 3.2 of the Supplementary Files list the required conditions for a match to be considered valid. Corroborating previous results, the fraction of pBLAST matches is also correlated with the quality of the transcript support level (Supplementary Figure A8–9).

#### 3.2.2 Input attribution score analysis

Several techniques exist that allow us to gain insights into the decision-making process of the trained model. Here, we apply integrated gradients to evaluate the relative contribution of the input nucleotides on the transcript to the predicted output. Integrated gradients, first introduced by Sundararajan et al. (2017), utilizes the partial derivatives of the model prediction with respect to the input values in order to assign attribution scores. We observe several expected patterns, such as the high importance of the candidate TIS itself and surrounding areas (e.g. Kozak sequence context). In addition to this, we see recurring patterns that both the translation termination sites and reading frame can have importance towards determining TISs (Figure 3b). No annotations on reading frames or translation termination sites were given to the model, meaning that biological features of translation were successfully learned automatically. We hypothesise that the relatively higher importance of the second and third element of the reading frame can be ascribed to their higher information content. Further analysis of these attribution score profiles might reveal additional biological factors influencing the translation process(Figure 3a).

#### 3.2.3 Detection of small proteins and multi-TIS transcripts

In 1994, it was postulated that the minimum length of functional proteins is around 100 amino acids (Dujon et al. 1994). Today, multiple studies have reported proteins shorter than 100 amino acids (Andrews and Rothnagel 2014; Vitorino et al. 2021), fulfilling roles in different types of regulatory mechanisms (Jorgensen and Dorantes-Acosta 2012; Ye et al. 2020). Nonetheless, small open reading frame encoded peptides (SEPs) continue to be underrepresented in existing annotations (Frith et al. 2006). With sequence alignment algorithms suffering from low statistical power for shorter sequences, more evidence is needed to differentiate the false from the true positives (Pauli et al. 2015).

The introduction of high-performing machine learning models could offer a solution to the detection of SEPs. The ability of the model to detect SEPs is reflected by the overlap between model predictions and a recently published list of newly introduced SEPs that were discovered using ribosome profiling (Mudge, Ruiz-Orera, Prensner, Brunet, Calvet, et al. 2022) (see Supplementary Table A6). Nonetheless, the absence of SEPs in the data used to train and validate novel approaches is likely to influence the model and evaluation process. We observe overall lower probability scores for TIS positions that result in smaller CDSs (Supplementary Figure A11).

Figure 4 shows some examples on the output of the TIS Transformer for several high-profile and validated SEPs. Akimoto et al. (2013) prove the existence of three upstream open reading frames (uORFs) for the MKKS gene through proteome analysis, serving as a regulatory mechanism (peptoswitch). uMKKS0, uMKKS1 and uMKKS2 are reported to be 43, 63, and 50 amino acids long, respectively (Figure 4a). Matsumoto et al. (2017) could validate the expression of a 90 amino acid sORF on the SPAAR gene (Figure 4b). The micropeptide is shown to be an important factor in regulating biological pathways related to muscle regeneration. Two studies reported the existence of a 56 SEP, found to affect mitochondrial respiration (Makarewich et al. 2018; Stein et al. 2018) (Figure 4c). Young and Wek (2016) report the existence of an upstream CDS of 53 codons overlapping with the ATF-4 coding region (Figure 4d). Supplementary Table A6 feature more information on model predictions of recently recovered small ORFs (Mudge, Ruiz-Orera, Prensner, Brunet, Gonzalez, et al. 2021). Supplementary Figure A12 gives the model output of transcripts with multiple high-ranking CDSs, some of which being short CDSs.

In contrast to the Ensembl database used for training, the model allows for multiple high ranking TIS per transcript (Figure 4a/b/d) and does not seem to be affected by the lack of multiple TISs in the training data. When selecting as many TISs within the positive set as are present in the Ensembl annotations (i.e. by incoporating the top ranking predictions in the positive set), a total of 956 transcripts with multiple TISs would be obtained. Supplementary Figure A10 shows the occurrence of upstream (overlapping), downstream (overlapping), and internal ORFs on the transcriptome annotated by TIS Transformer. Several examples of such transcripts are shown in Supplementary Figure A12.

## 4 Discussion

Recent advancements of machine learning in processing sequential data, mainly introduced in the field of natural language processing, portend new opportunities towards the creation of predictive models on biological sequence data. The introduction of the FAVOR+ mechanism, which reduces the complexity of the attention step, has been an essential advancement to make processing of full transcript sequences at single nucleotide resolution possible. In this study, we investigate the use of these transformer networks to determine TIS sites based on the processed transcript sequence. We introduce TIS Transformer and benchmark it with previous solutions and show that a major improvement in performance was achieved. Transformers offer several advantages, notably, the model is able to parse variable length inputs, allowing it to process the full transcript sequence. It is efficient in parsing transcript information, as it requires only to parse the full transcript once to provide predictions for all its nucleotide positions. Furthermore, we see that improved performances have been achieved with models containing a comparable number of trained weights.

Although the research objective has been taken on by a plethora of studies in the past, the general usability and advantage of utilizing machine learning for annotating the full transcriptome has remained unclear. Various factors that previously posed as limitations, such as computational requirements, do not necessarily prove to be an inhibiting factor today. Rather, limitations in performances are indicated to be caused by faulty transcript sequences or noisy TISs annotations, as opposed to shortcomings within the proposed model architecture. This hypothesis is further substantiated when evaluating the fraction of matches returned by pBLAST on novel TISs predictions. The positive correlation with the model certainty speaks to the ability of the model to find novel TIS sites (see Figure 2a, while the negative correlation with the transcript support level (see Supplementary Figure A8–9) speaks to the potential flaws within the transcriptome mapping. Thus, future work will need to be focused on both improving the mapping of the transcriptome and proteome.

In this study, the positive set incorporates all annotations provided by Ensembl, thereby incorporating a set of transcript types that impede the translation process. For example, included are protein products starting from TISs that eventually are broken down in the nonsense mediated-decay process and products originating from transcripts missing validated coding mRNA ends (GENCODE tags CDS_end_NF/mRNA_end_NF). A large portion of novel annotations performed by the model occurs on special type transcripts (see Supplementary Figure A10). Future work might focus on differentiating between types of translation events, such as those resulting in nonsense mediated decay, as it is not clear how the inclusion of these groups influence the model predictions.

Evaluation of the decision-making process of the model has corroborated the ability of deep learning to achieve automated feature extraction. Patterns can be observed that match properties of translation, where the reading frames, TISs, and TTSs can be discerned from attribution profiles (Figure 3). The impact of biases introduced by existing (flawed) data sets is not clear. Biases are expected to be present within the positive set, such as the lack of TISs that result in SEPs. Other problematic TISs could include those that result in unstable proteins as these are harder to pinpoint using *in vivo* detection methods Ruiz Cuevas et al. (2021). It is possible that the importance of the amino acid compositions or the length of the protein, and thereby the location of the TTS, is a direct result of these existing biases. Another example is the absence of near-cognate initiation codons from the training data and predictions, with only 7 (0.006%) instances (all CTG) present in in the top (k) predictions of all chromosomes.

The relevance of deep learning has strongly increased in recent years as its application and adaptation becomes more widespread. Most notably has been the release of AlphaFold, which has become a central tool for protein structure prediction (Jumper et al. 2021). Similarly, annotation software driven by machine learning, such as TIS transformer, can drive the design of future studies or serve as an extra validation step. These can be focused on the discovery of new proteins, improving our understanding on the biological drivers of translation, or predicting the occurrence of TISs on novel transcripts, an required step in the study of biological pathways.

## 5 Data availability

The main repository is available at https://github.com/jdcla/TIS_transformer. All annotations of the model are publicly available. All code is available and can be used to train varying model architectures or remap the proteome of other organisms. All discussed models are made available for the community, and can thereby be applied to custom transcripts. The input data, unprocessed model outputs, and curated predictions for each chromosome, as used in this study, are available. Lastly, to promote access to the results, and allow users to quickly obtain predicted TISs given certain criteria, we provide an online tool that is linked on the GitHub and currently online at https://jdcla.ugent.be/TIS_transformer. To illustrate, one can easily collect the small ORFs on non-coding sequences, all transcripts featuring multiple TISs, and transcripts featuring upstream ORFs.

## 6 Funding

The work presented was sponsored by Novo Nordisk Research Centre Oxford Ltd with the work being carried out jointly by Ghent University and Novo Nordisk employees. Novo Nordisk took part in the study design and overall supervision and guidance of the project, with a focus on the evaluation of biological relevance. Ghent University was responsible for the study design, technical development, testing and implementation of the deep learning model.

## Supplementary Information and Figures

### 1. Model architecture

The model architecture consists of multiple layers that are identical in structure but feature unique trainable model parameters. Two sets of embeddings are used; trainable nucleotide embeddings (A, C, G, T, N, [START], [STOP], [MASK]) and fixed positional embeddings. The transformer structure features multiple layers with multiple attention heads per layer. The workings of the Performer network are described by Choromanski et al. [2021]. The outputs of the network are send to a set of fully connected layers to obtain a binary output at each input position.

#### Algorithm 1 TIS Transformer network architecture. Given are the different layers, their respective dimensions as defined by their hyperparameter names, the dimensions for TIS Transformer (Table A4), and the resulting total weights.

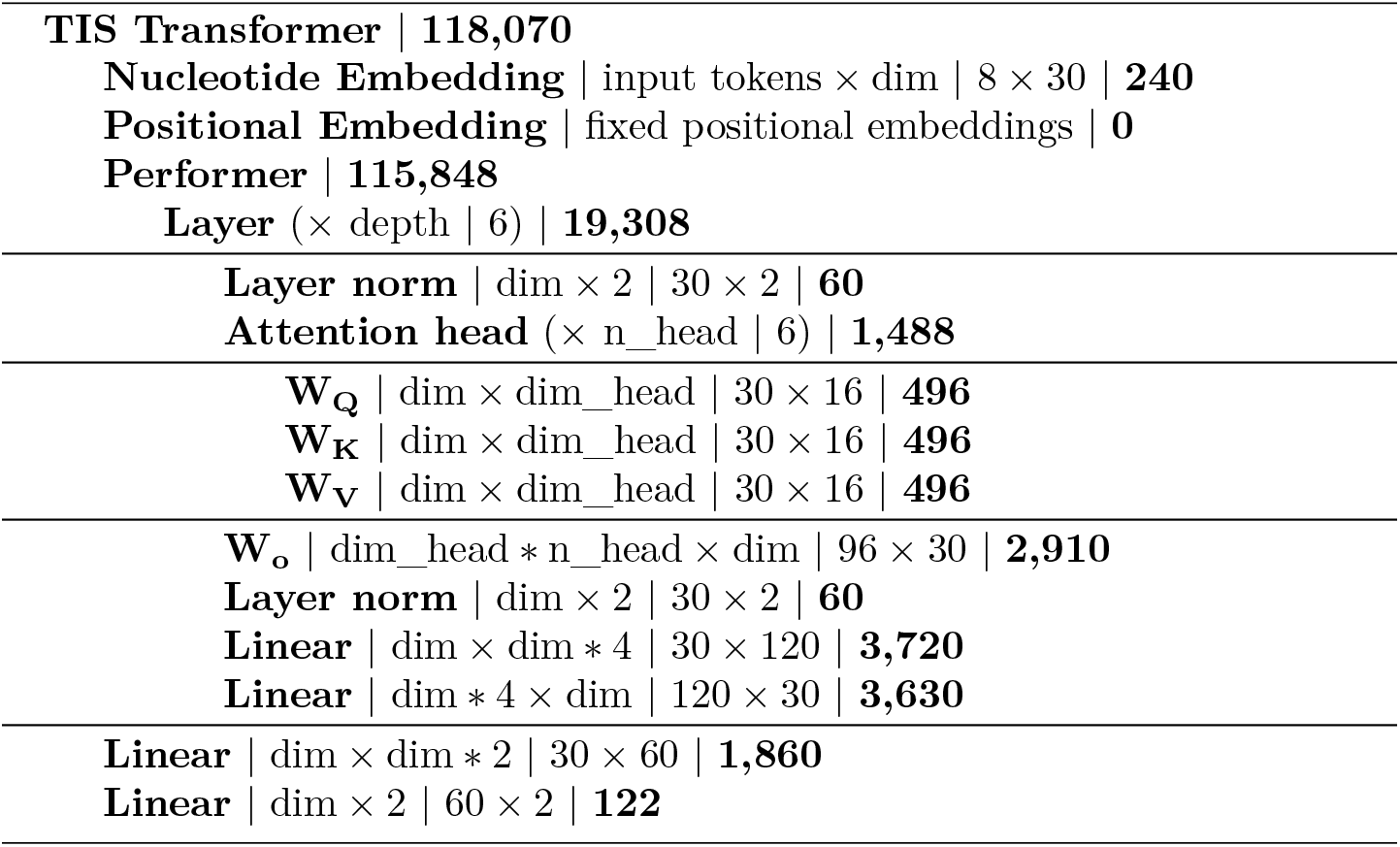

### 2. Model selection

#### 2.1. Hyperparameter optimization

Hyperparameter optimization is performed for a single set-up: transcripts from chromosomes 1, 7, 13, and 19 are excluded (test set) and chromosomes 2 and 14 are applied to select the optimal hyperparameters (validation set). Overall, no individual hyperparameters were observed to be more effective than others in improving performances. It was observed that a correlation exists between the total number of model parameters and model performance. To reflect this, the performances of three model architectures are given; TIS Transformer S(mall), TIS Transformer and TIS Transformer (L)arge (see Table A2, A3, A4, A5). Each network represents a tripling of model parameters. Performance gains showed to be most substantial when increasing the model parameters (i.e. number of layers, number and dimensions of attention heads, dimension of the hidden state) up to a certain point, after which gains stagnate. Although three times bigger, the performance of the TIS Transformer L is marginally better than that of TIS Transformer. The minimal loss on the validation set for both architectures is similar before they both start overfitting (see Figure A4). These findings reflect those given in the main manuscript, where further improvements of machine learning approaches are likely to be hampered by a set of noisy annotations. This was shown through the correlation of performances with the support level of the transcripts and the verification of other annotation platforms such as CCDS.

Notwithstanding the size of the data set and overall high computational requirements of transformer architectures, model optimization is possible on a single RTX 3090 and converged after ca. 10 hours due to the relative shallowness of the final transformer architecture (Supplementary Tables A3–A5). The details of varying model architecture performances are given by Supplementary Table A2 and Supplementary Figure A4, A5, A6.

**Table A1:** Overview of studies on TIS annotation using sequence information. For every study is given, the year of publication, the name of the tool, the machine learning approach (SVM: Support Vector Machine, NN: Neural Network), the size of the input sequence around the candidate TIS, the total number of model parameters, the total number of data samples used in the study, and the location of public code repository. denotes that the value does not apply. ‘?’ is used when the answer is unclear from the manuscript.

| Author | Year | Model name | Type | Input size | # parameters | # Samples | Code Repo |
| --- | --- | --- | --- | --- | --- | --- | --- |
| Zien et al. [2000] | 2000 | - | SVM | 200nt | Variable | 13,503 | - |
| Saeys et al. [2007] | 2007 | StartScan | Varia | 200nt | - | 1,267,701 | - |
| Chen et al. [2014] | 2014 | iTIS-PseTNC | SVM | 198nt | Variable | 2,318 | - |
| Kabir et al. [2015] | 2015 | iTIS-PseKNC | SVM | 198nt | Variable | 2,318 | - |
| Goel et al. [2020] | 2020 | - | SVM | 303nt | Variable | 3,020 | - |
| Zhang et al. [2017] | 2017 | TITER | NN | 203nt | ~431K | 104,675 | GitHub |
| Zuallaert et al. [2018] | 2018 | TISRover | NN | 203nt | ~240K | 94,642 | - |
| Kalkatawi et al. [2019] | 2019 | DeepGSR | NN | 203nt | ~181M | 28,244 | Zenodo |
| Wei et al. [2021] | 2021 | DeepTIS | NN | ? | ? | 115,728 | GitHub |
| Clauwaert et al. | 2022 | TIS Transf. | NN | Variable | ~118K/356K | 431,011,438 | GitHub |

#### 2.2. Benchmarking

A multitude of studies have previously been performed applying machine learning techniques for the prediction of TISs. Previous studies utilizing the transcript nucleotide sequence have been listed in Table A1 Zien et al. [2000], Saeys et al. [2007], Chen et al. [2014], Kabir et al. [2015], Goel et al. [2020], Zhang et al. [2017], Zuallaert et al. [2018], Kalkatawi et al. [2019], Wei et al. [2021]. In contrast to existence of multiple studies, no single data set exists that functions as the go-to benchmark data set for this problem setting. This can be attributed to various reasons, the main ones being the computational limitations of some techniques making it impossible to process multiple millions of samples, and the obscurity of the ground truth, resulting in multiple existing platforms each featuring varying sets of annotated TISs.

In this study, we utilize the full genome of the selected organism (Homo sapiens) of Ensembl to train, validate, and test the model with. We believe this approach to have several advantages over custom subsampled data sets. The vast majority of transcript positions are non-TIS sites, where sub-sampling mainly affects the negative set. The technique used to sub-sample the negative set influences the population sampled, and thus the resulting performances. To illustrate, a model sampling ca. 10,000 samples of the negative set at random effectively covers only 0.002% of the population. For a setting where 0.01% of the negative samples bear sequence similarity to the region of an actual TIS position (i.e. hard to predict), this would result in five such samples in the negative set. Performances measured on such models will easily result in near-perfect precision and accuracy scores. However, it fails to portray the model’s capability when applied on the full transcriptome, where the vast number of negative samples results in a set of false positives that heavily outweighs the number of positive samples. Most studies aim to balance the number of positive and negative samples. While several studies discuss an approach that seeks to sample the negative samples that have similarity with the positive samples (i.e. hard to predict), each study follows a new approach. As such, variations between various methods of sampling causes resulting model performances to vary. This is illustrated by our results contradicting published results. TITER performs better than DeepGSR as published by [Kalkatawi et al., 2019].

It is impossible to perform a benchmark against approaches utilizing support vector machines due to the size of the data. Large data sets are required to train for neural networks, but pose a problem for support vector machines, where the training and evaluation time scale quadratically with the number of samples processed. It is nonetheless implausible that support vector machines can offer comparable performances considering the low number of samples they are trained with. Additionally, all previous studies incorporating support vector machines as part of their benchmark list these models as inferior to neural network implementations [Zuallaert et al., 2018, Wei et al., 2021, Kalkatawi et al., 2019]. Lastly, we were unable to apply DeepTIS [Wei et al., 2021] due to a lack of information given in the paper. Some of the missing or unclear details include specific hyperparameter values (e.g. *q* which defines the input window length) or the exact model architecture of ’DeepTIS2’, featured in the paper as the best performing one. The online GitHub repository seemed incomplete as there was no code utilizing recurrent neural networks, which should be part of the backbone of ’DeepTIS2’. We were unable to reach the authors of this work for clarification.

To ensure a fair comparison of the listed methods, all methods are being trained and evaluated on the exact same data sets. Due to several constraints imposed by individual methods, all transcript positions matching at least one of these constraints are excluded from the data. These constraints are: only ATG sites, only positions on transcripts with a length of less than 30,000 nucleotides, only transcript positions that are distanced at least 300 nucleotides from the start and end of the transcript, and no ’N’ annotated nucleotides within a 300nt window of the candidate TIS site. Applying these constraints on the full results in a train, validation and test set of 3,608,307, 641,264 and 1,069,321 candidate TIS positions. Due to several available network architectures being implemented using outdated software packages for GPU accelerated computation (TISRover: Lasagne, TITER: Theano), the decision was made to re-implement all models using PyTorch (Lightning). Since convergence of the TITER model took ca. 24 hours to complete, we decided to forego the training and use of 32 individual models with which a prediction is made due to computational requirements. The use of 32 independent neural networks is cited to have further improved results by Zhang et al. [2017], likely due to reduction of the variance error. Nonetheless, it is clear that this step would not close the performance gap between a single TITER model and the TIS Transformer model. All scripts used to perform the benchmark are found in the public GitHub repository https://github.com/jdcla/TIS_transformer/tree/main/benchmarks. By cross-referencing the total number of model weights we have verified the correct implementation of each network architecture.

### 3. Results analysis

#### 3.1. Rank (k) and ’false positives’

With hundreds of millions of predictions, it is necessary to select only a subset of predictions for further analysis. Following the total number of Ensembl TIS annotations *k* unique to each chromosome, we have determined a custom ranking for the predictions of each chromosome that is scaled to this number (rank k). For a chromosome with 1000 positively annotated TIS, the highest ranking output gets rank (k): 0, the 500th highest output rank(k): 0.5, and the 2000th highest prediction a rank (k): 2. Applying this ranking, it is possible to get a quick idea on how the model prediction compares to other predictions within the chromosome. In general, false positives refer to positive predictions with rank (k) < 1 that were not previously annotated by Ensembl.

#### 3.2. pBLAST

To cross-reference model predictions that are not featured by Ensembl, we evaluated the CDSs resulting from predicted TISs using pBLAST. These were queried against Swiss-Prot, TrEMBL (mammals) and all supplementary isoforms. pBLAST matches were filtered following three constraints based on various properties: a match requires to be at least 95% identical to the query sequence, a maximum difference in length between the query and match of 5%, and a maximum difference in distance between the aligned start and stop sites of 5% of their total length. We are aware that given constraints are not perfect. However, constraints were tuned by evaluating the total number of matches returned when evaluating the pBLAST results on the annotations provided by Ensembl, and found to return a large portion of the correct proteins without including some obvious false positives.

#### 3.3. Online result browser

In addition to featuring the raw results, the code and the scripts used to obtain the results on the public GitHub repository, we host an online server that provides a more accessible approach towards making our findings open to the public. The tool is accessible through https://jdcla.ugent.be/TIS_Transformer. The link is furthermore linked through our public GitHub repository at https://github.com/jdcla/TIS_transformer. With this, we hope to attract a larger group of users that is otherwise not experienced with coding or data manipulation.

The result browser allows the user to filter predictions based on various features. Currently implemented filters are: gene/transcript name, ORF type, ORF length, Ensembl annotation, transcript type, prediction rank, and number of matches on transcript. The query returns a table of all matches, and all related information of the TISs and resulting CDSs, for easy download and visualisation. To illustrate, one can easily collect the small ORFs on non-coding sequences, all transcripts featuring multiple TISs, and transcripts featuring upstream ORFs.

**Figure A1:**
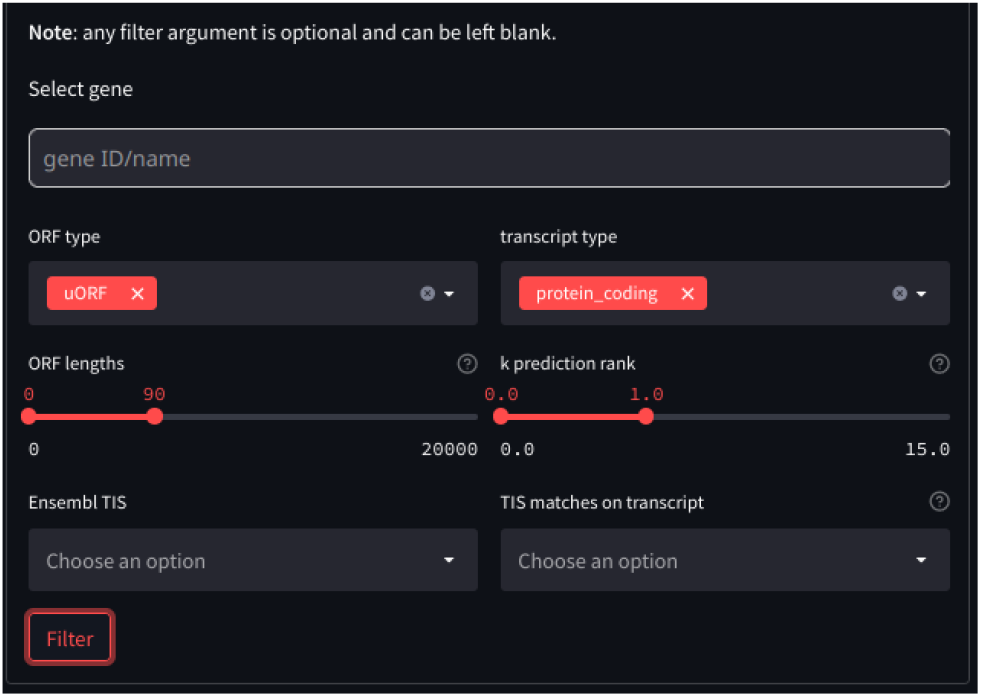
A screenshot of the result browser at https://jdcla.ugent.be/TIS_transformer featuring multiple filter arguments.

**Figure A2:**
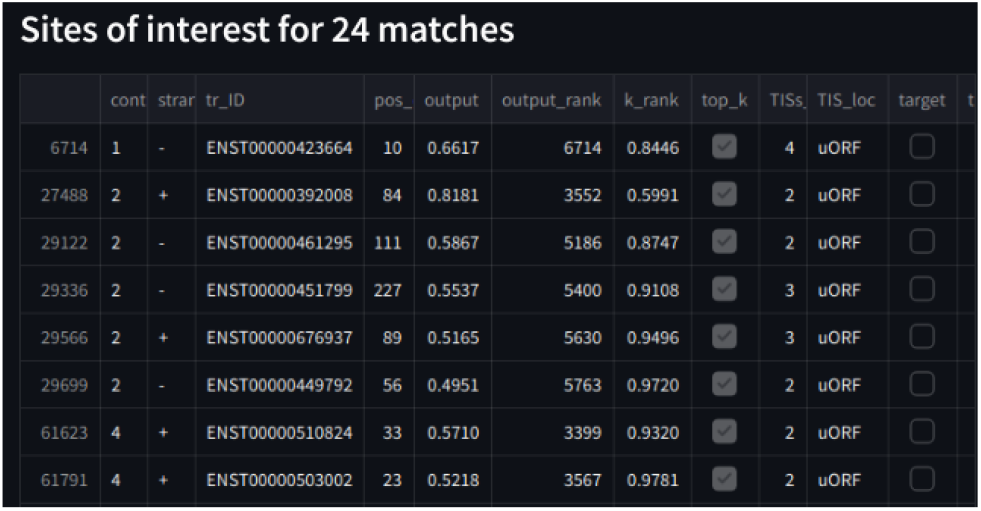
A screenshot of part of the resulting table at https://jdcla.ugent.be/TIS_transformer after filtering results as featured in Figure A1.

**Figure A3:**
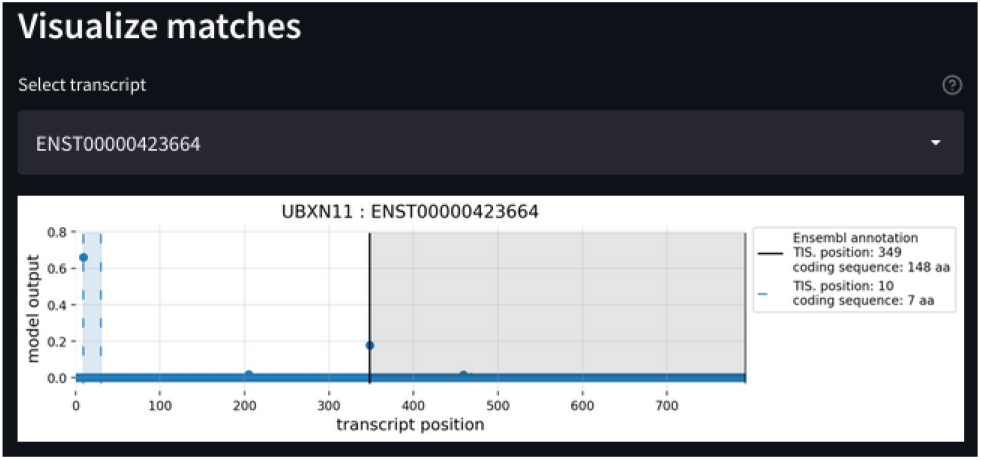
A screenshot of a visualization offered at https://jdcla.ugent.be/TIS_transformer of one of the predicted TIS that matches the filter arguments given in Figure A1.

### 4. Supplementary Tables

**Table A2:**
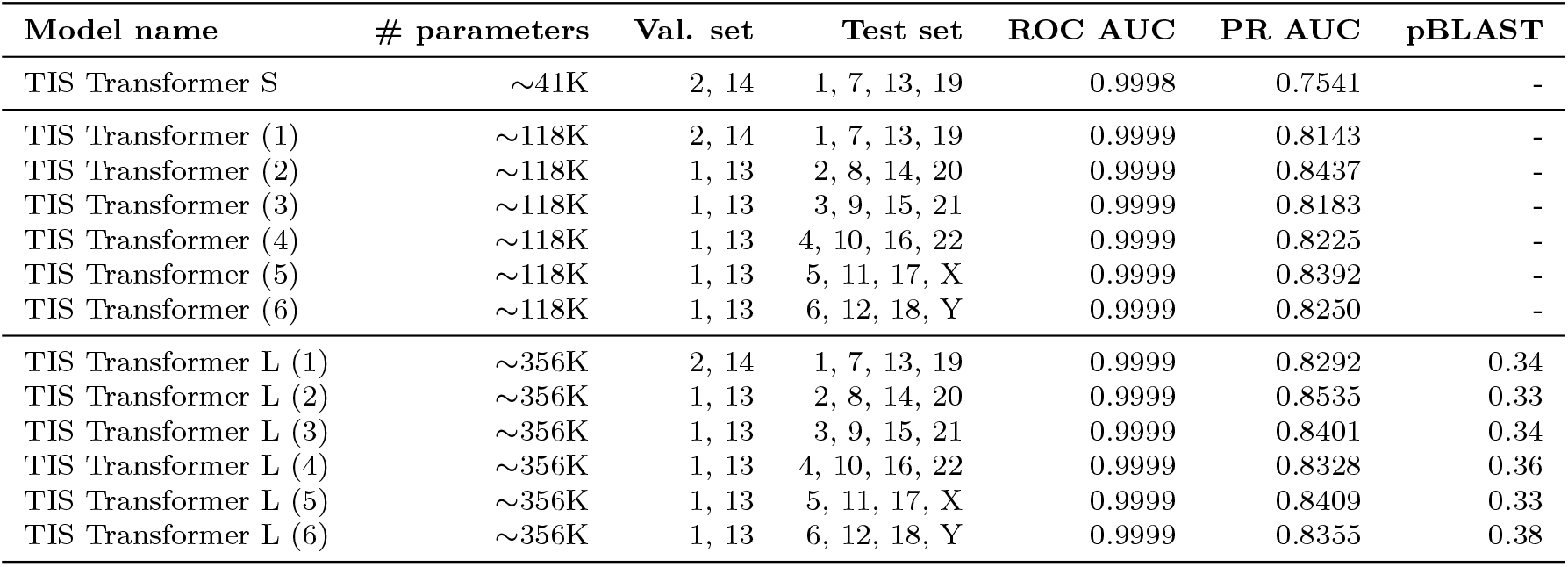
Model performances for the varying models used to remap the human proteome. The test and validation sets refer the the contig identifiers. The training set uses all remaining contigs. pBLAST refers to the fraction of false positives TISs (rank (k) < 1) that return a match when performing pBLAST search on their resulting CDSs.

**Table A3:** Overview of the hyperparameters that define the TIS Transformer S(mall) model architecture. This model was used as a step to compare different architectures. Also given are the keys used to define the model using the code at https://github.com/jdcla/TIS_transformer

| Hyperparameter | Argument | Value | Hyperparameter | Argument | Value |
| --- | --- | --- | --- | --- | --- |
| learning rate | -lr | 0.001 | dim. of heads | -dim_head |  |
| dim. of hidden state | -dim | 20 | kernel function | -kernel_fn | torch.nn.ReLU() |
| layers | -depth | 4 | local window size | -local_window_size | 256 |
| attention heads (layer) | -heads | 4 | local attention heads | -local_attn_heads | 3 |
| Total trainable model parameters |  | ~41K |  |  |  |

**Table A4:** Overview of the hyperparameters that define the TIS Transformer model architecture. Also given are the keys used to define the model using the code at https://github.com/jdcla/TIS_transformer

| Hyperparameter | argument | value | Hyperparameter | argument | value |
| --- | --- | --- | --- | --- | --- |
| learning rate | -lr | 0.001 | dim. of heads | -dim_head | 16 |
| dim. of hidden state | -dim | 30 | kernel function | -kernel_fn | torch.nn.ReLU() |
| layers | -depth | 6 | local window size | -local_window_size | 256 |
| attention heads (layer) | -heads | 6 | local attention heads | -local_attn_heads | 4 |
| Total trainable model parameters |  | ~118K |  |  |  |

**Table A5:** Overview of the hyperparameters that define the TIS Transformer L(arge) model architecture. This model was used as a step to compare different architectures. Also given are the keys used to define the model using the code at https://github.com/jdcla/TIS_transformer

| Hyperparameter | Argument | Value | Hyperparameter | Argument | Value |
| --- | --- | --- | --- | --- | --- |
| learning rate | -lr | 0.001 | dim. of heads | -dim_head | 16 |
| dim. of hidden state | -dim | 48 | kernel function | -kernel_fn | torch.nn.ReLU() |
| layers | -depth | 8 | local window size | -local_window_size | 256 |
| attention heads (layer) | -heads | 8 | local attention heads | -local_attn_heads | 5 |
| Total trainable model parameters |  | ~356K |  |  |  |

**Table A6:**
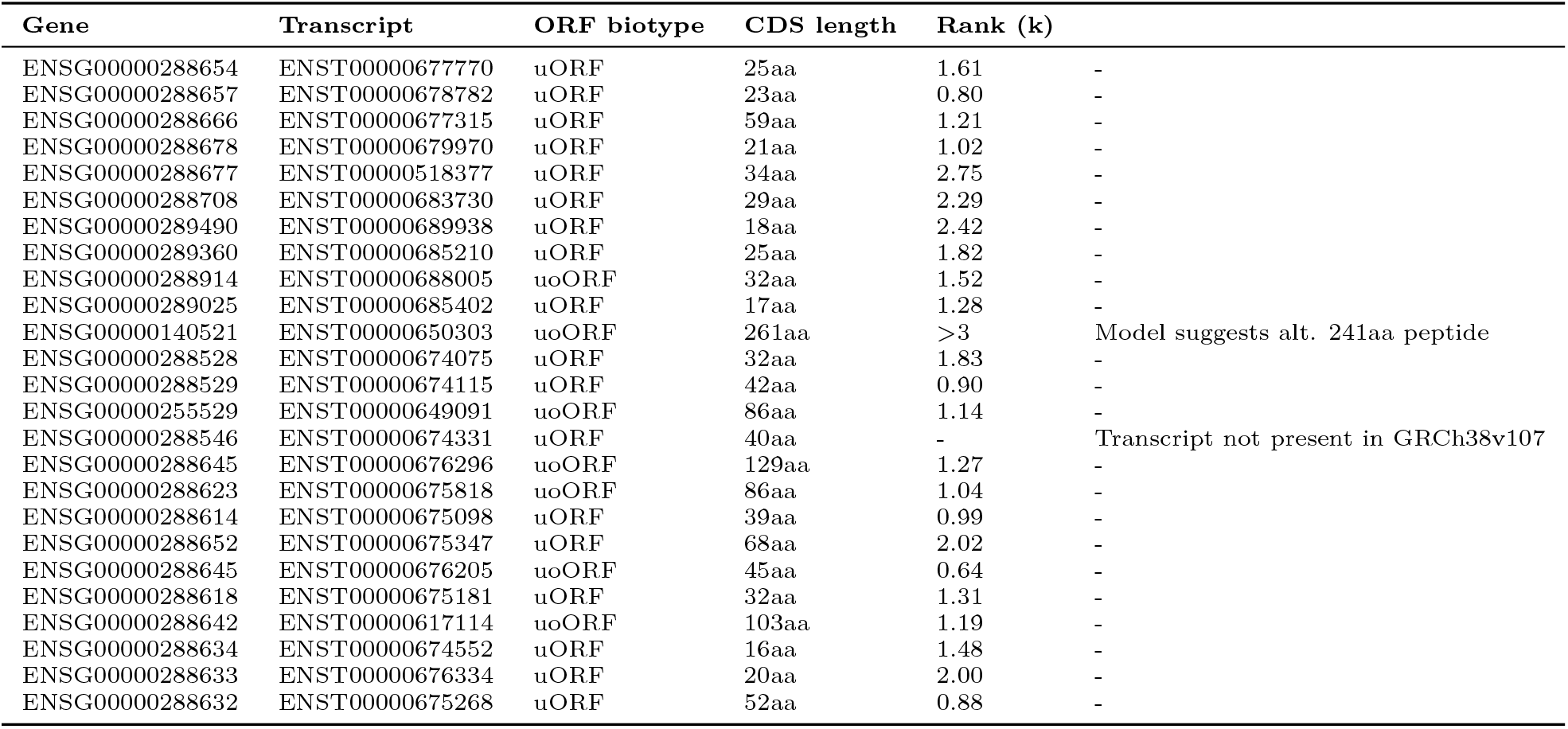
A set of ORF annotated sequences that have been recently added to GENCODE as part of Ribo-seq studies. The list has been retrieved from a recent publication on the advancement of ORF detection through a community-led framework Mudge et al. [2022]. uORF: upstream open reading frame; uoORF: upstream overlapping open reading frame.

### 5. Supplementary Figures

**Figure A4:**
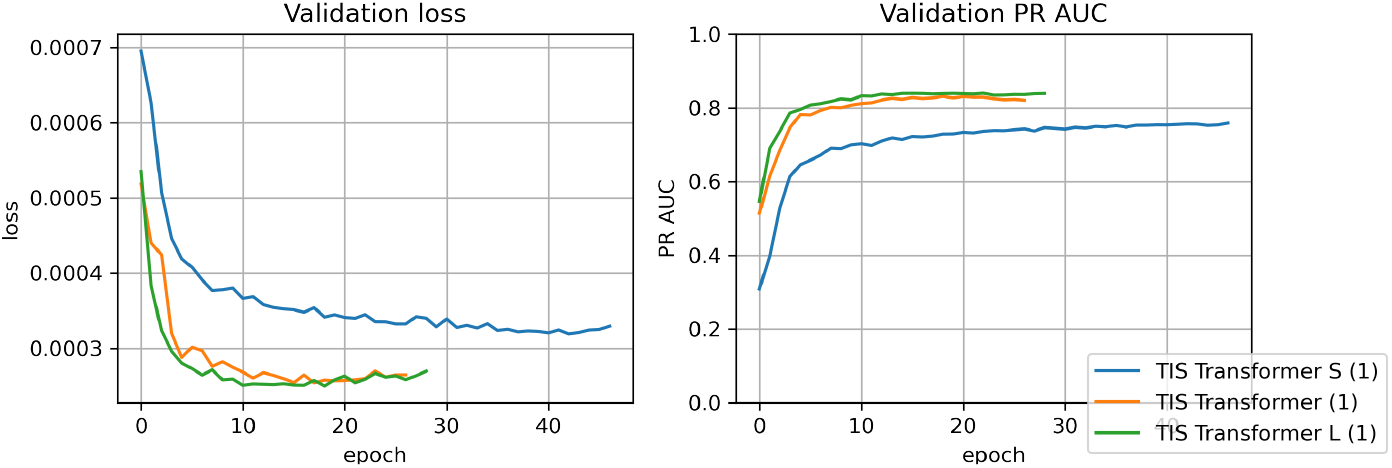
The loss and PR AUC curves of three model architectures trained for annotating TISs. The validation and test sets used are chromosomes 2, 14 and chromosomes 1, 7, 13, 19, respectively. The hyperparameters for each model are given in Table A3, A4, A5

**Figure A5:**
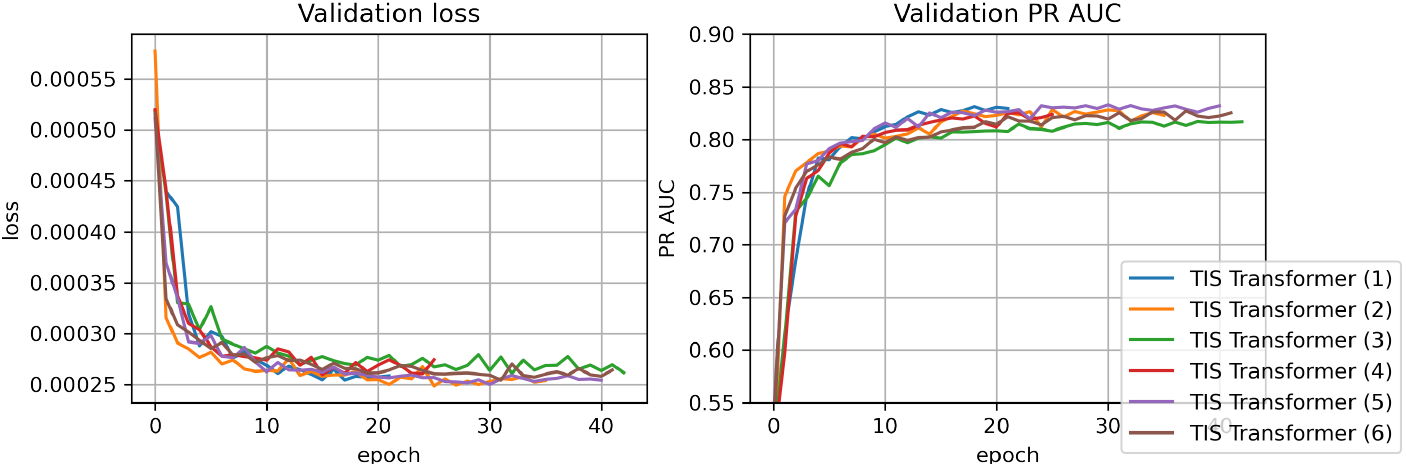
The loss and PR AUC curves for the models trained on the TIS annotation task. Each model has a different set of chromosomes for the train/test/validation set, as given in Table A2.

**Figure A6:**
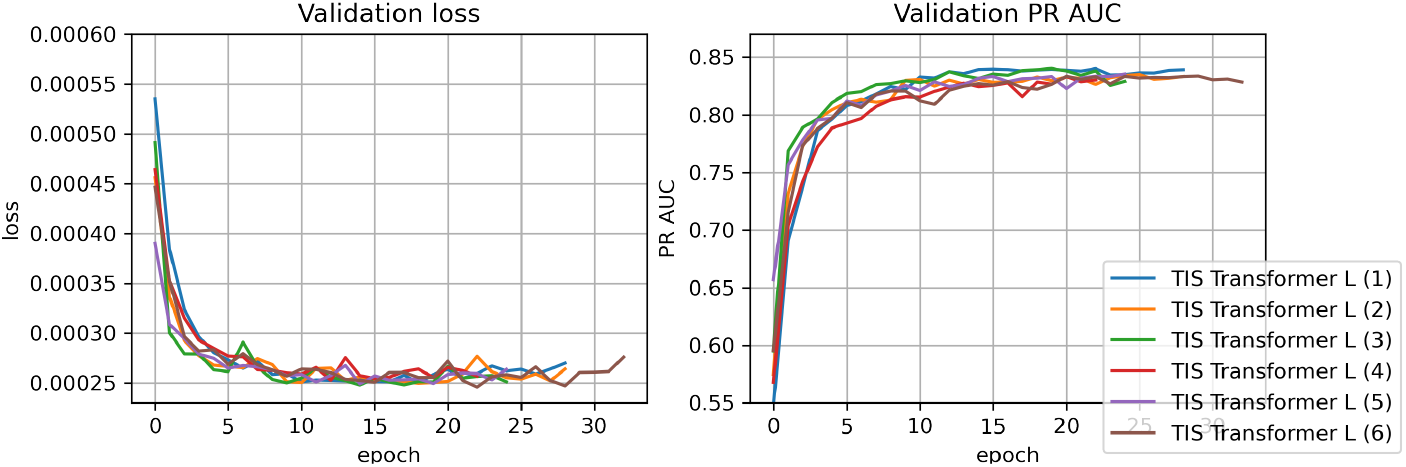
The loss and PR AUC curves for the models trained on the TIS annotation task. Each model has a different set of chromosomes for the train/test/validation set, as given in Table A2.

**Figure A7:**
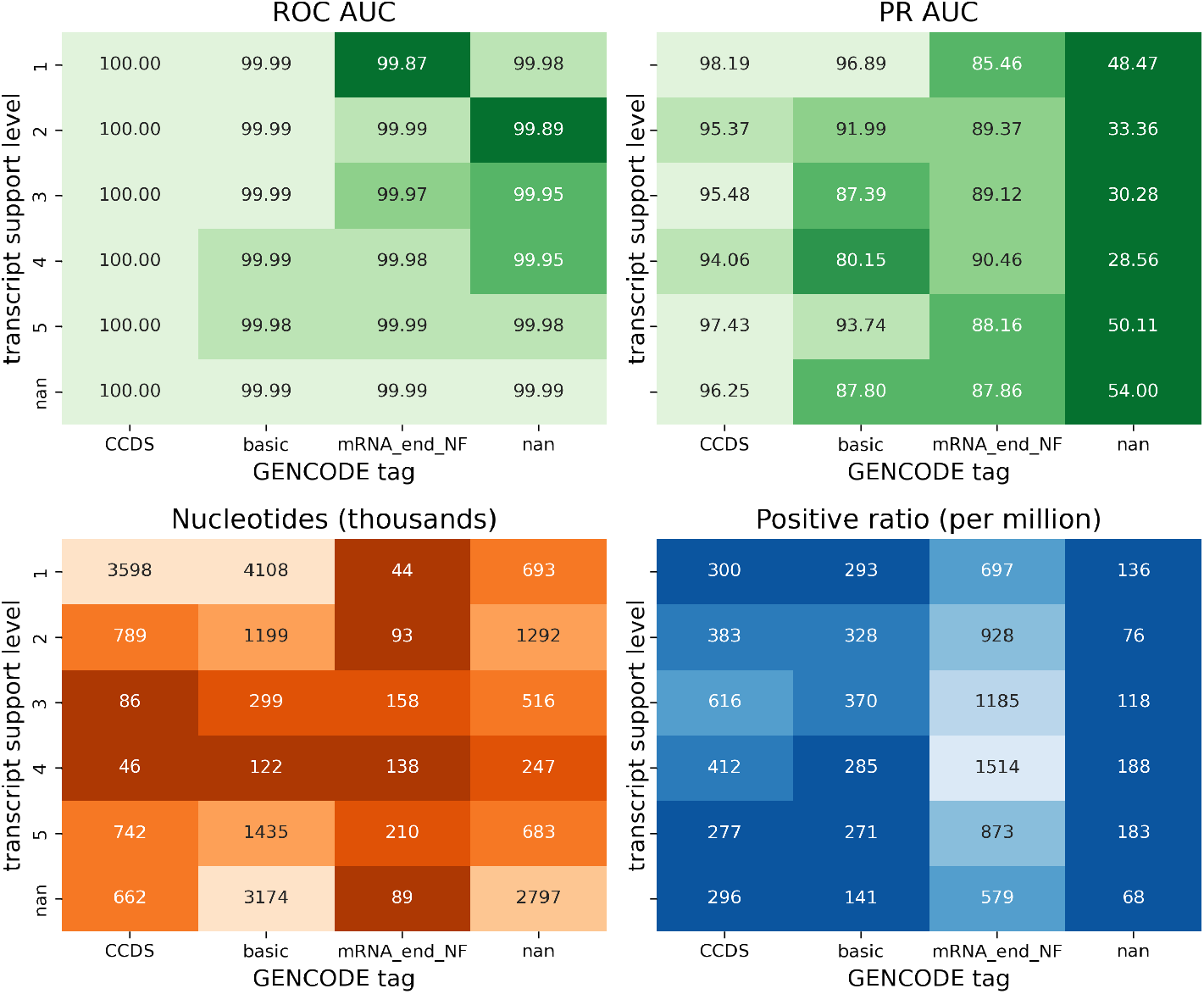
Model performances and input information for the inputs binned by transcript support level and tags given to the annotated translation initiation sites. ROC AUC and PR AUC performances (top) are given as well as the total number of annotated TISs (by Ensembl) and ratio of positive samples (w.r.t. negative samples). Values are obtained by binning predictions per transcripts according to transcript support level and by binning the predictions by tags given to the annotated translation initiation site.

**Figure A8:**
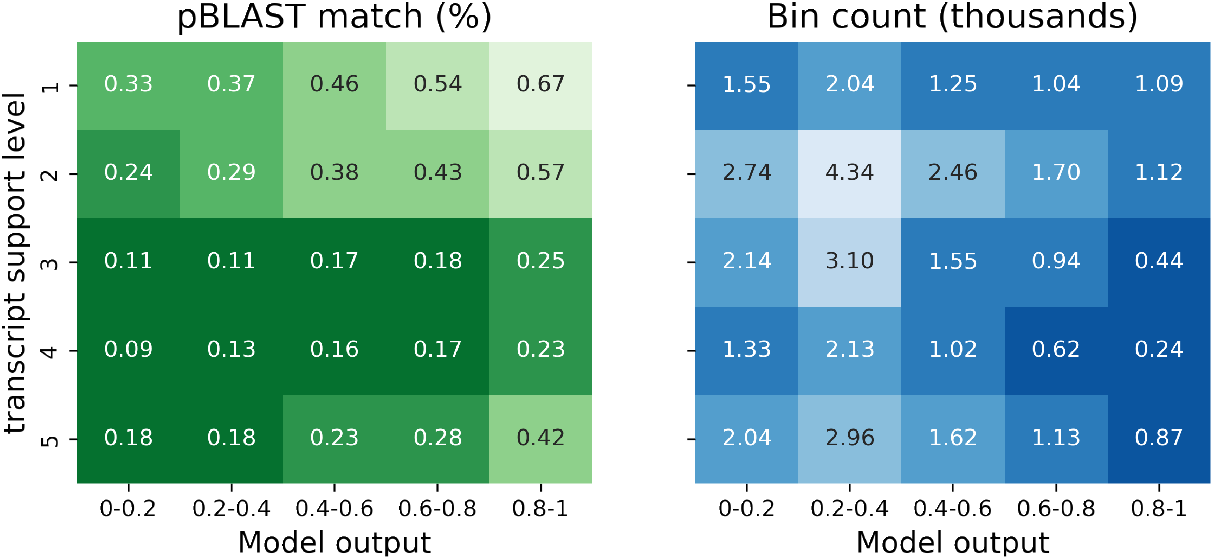
Fraction of good pBLAST matches for novel model predictions binned by transcript support level and model output range. Evaluated predictions have been limited to those within rank (k) < 1.5 (to equalize the leftmost bin size) and exclude those previously annotated by Ensembl. (**left**) The fraction of TISs that result in a coding sequence with a strong pBLAST match. (**right**) The number of samples in each bin.

**Figure A9:**
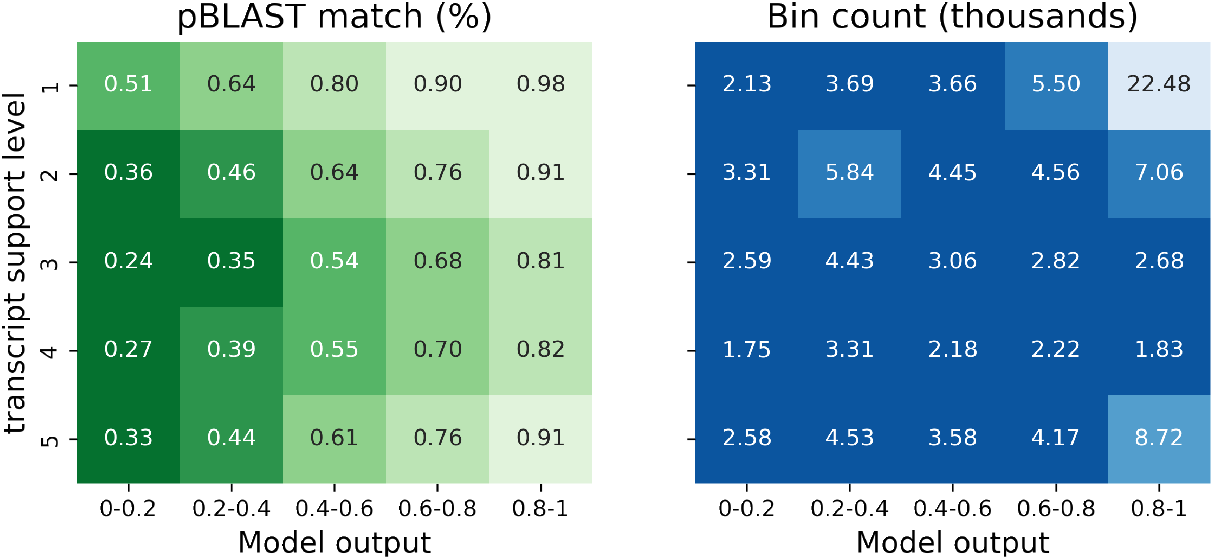
Fraction of good pBLAST matches for model predictions binned by transcript support level and model output range. Evaluated predictions have been limited to those within rank (k) < 1.5 (to equalize the leftmost bin size) and include those previously annotated by Ensembl. (**left**) The fraction of TISs that result in a coding sequence with a strong pBLAST match. (**right**) The number of samples in each bin.

**Figure A10:**
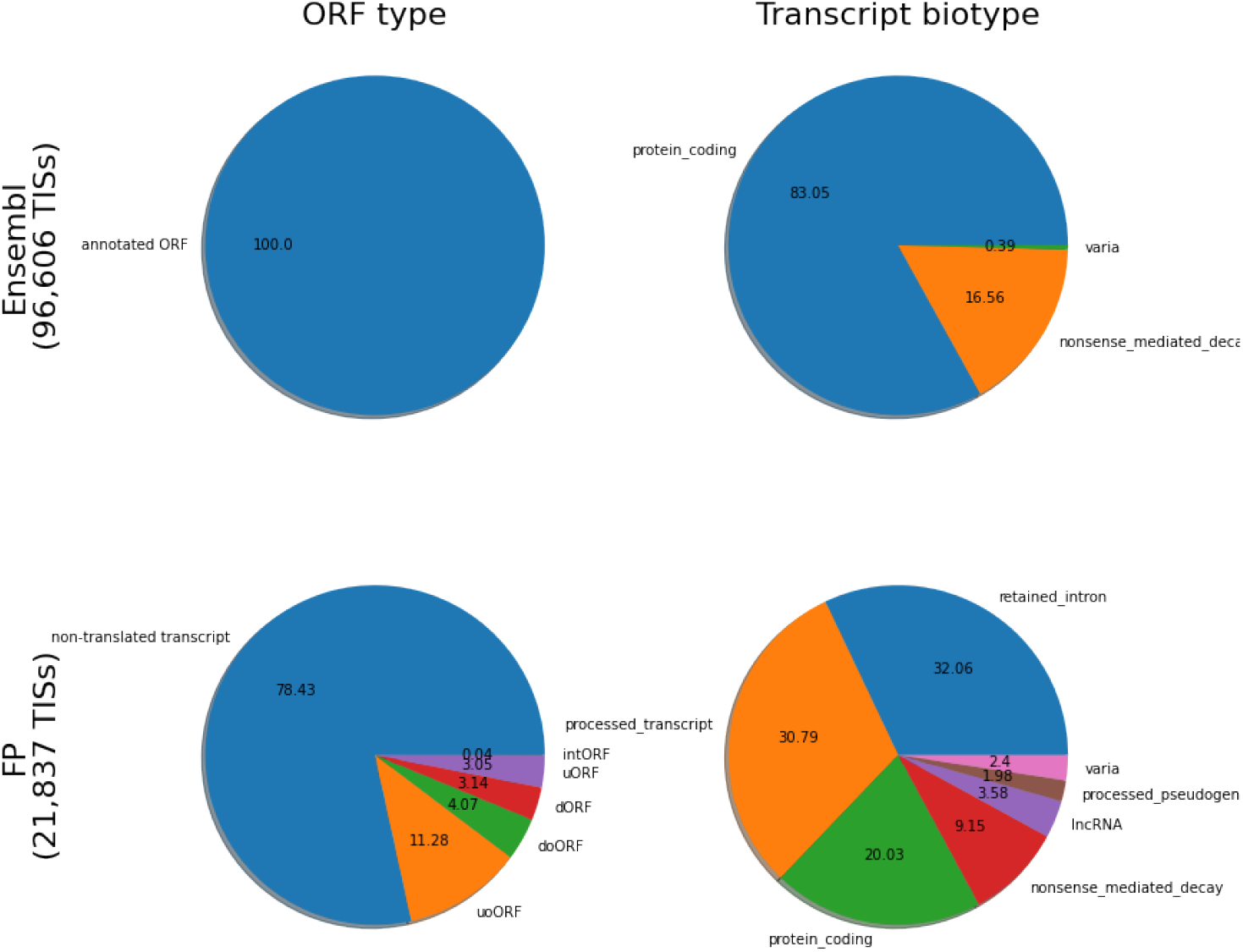
Property distributions between annotated and newly predicted TISs. These predictions constitute those that are not annotated by Ensembl and have a rank (k) < 1, referred to as the false positive (FP) set. For both groups the ORF type of the annotation and the biotype of the transcript are given. FP annotations on protein coding transcripts are either additional CDSs detected alongside the canonical annotation, or alternative TIS.

**Figure A11:**
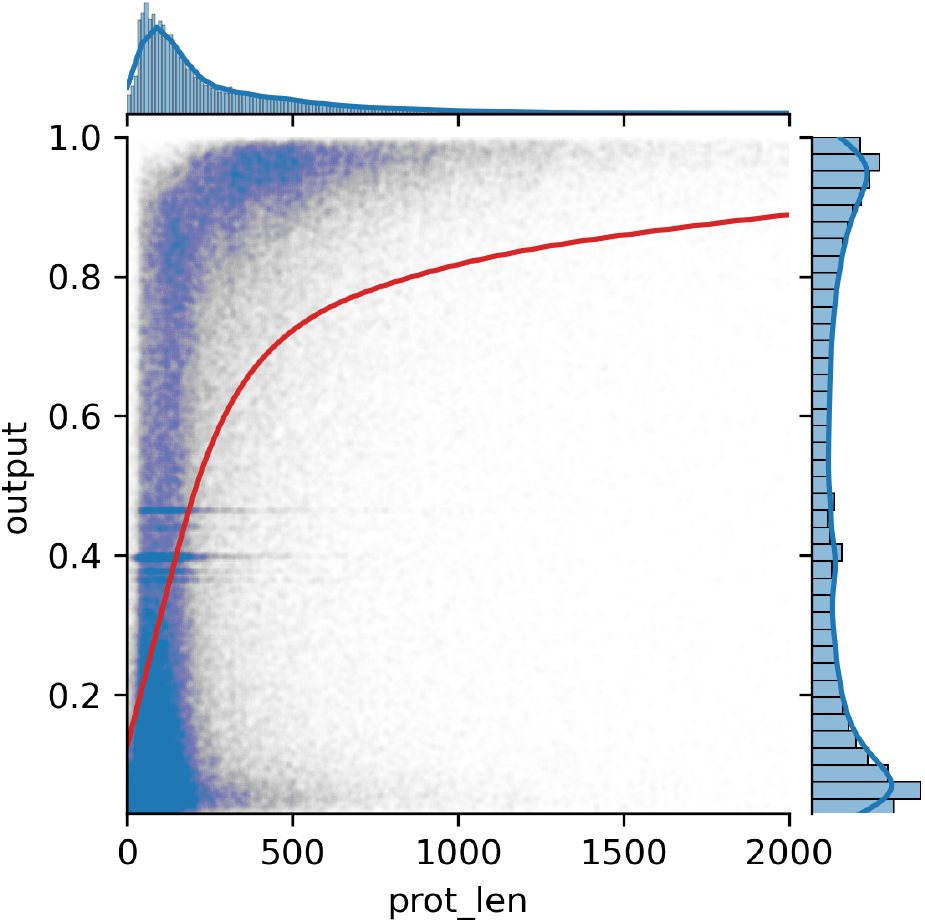
Correlation between the model output for a given translation initiation site and the length of its resulting protein. The trend line (red line) is obtained using LOWESS.

**Figure A12:**
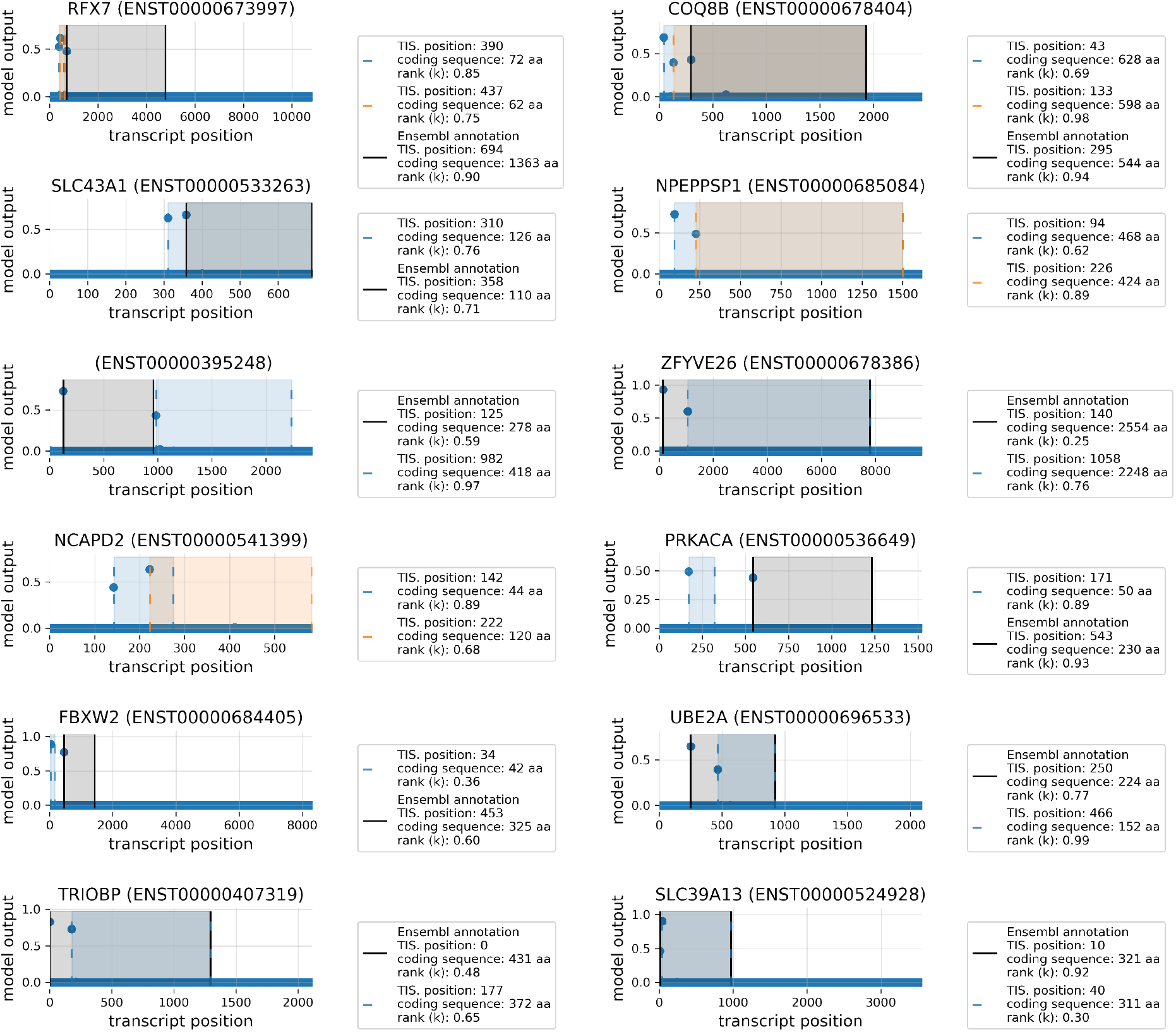
Model predictions on several transcripts with multiple high-ranking TIS positions. Shown are the model outputs (y-axis) for each position of the transcript (x-axis). For high TIS predictions, the bounds of the resulting CDS are given, as well as their length and prediction rank (*n*-highest prediction) on the chromosome.

## Notes

https://github.com/jdcla/TIS_transformer

### Competing Interest Statement

The authors have declared no competing interest.

### Summary of Updates

The manuscript has been overhauled using the updated Ensembl GRCh38 V107 (from v102).

## References

Aken BL et al. 2016. The Ensembl Gene Annotation System. Database. 2016:

Akimoto C, Sakashita E, Kasashima K, Kuroiwa K, Tominaga K, Hamamoto T, and Endo H. 2013. Translational Repression of the McKusickKaufman Syndrome Transcript by Unique Upstream Open Reading Frames Encoding Mitochondrial Proteins with Alternative Polyadenylation Sites. Biochimica et Biophysica Acta (BBA) - General Subjects. 1830: 2728–2738.

Andrews SJ and Rothnagel JA. 2014. Emerging Evidence for Functional Peptides Encoded by Short Open Reading Frames. Nature Reviews Genetics. 15: 193–204.

Chen W, Feng PM, Deng EZ, Lin H, and Chou KC. 2014. iTIS-PseTNC: A Sequence-Based Predictor for Identifying Translation Initiation Site in Human Genes Using Pseudo Trinucleotide Composition. Analytical Biochemistry. 462: 76–83.

Cheng J, Dong L, and Lapata M. 2016. Long Short-Term Memory-Networks for Machine Reading. arXiv:1601.06733 [cs].

Choromanski K et al. 2021. Rethinking Attention with Performers. arXiv:2009.14794 [cs, stat].

Dujon B et al. 1994. Complete DNA Sequence of Yeast Chromosome XI. Nature. 369: 371–378.

Eraslan G, Avsec, Gagneur J, and Theis FJ. 2019. Deep Learning: New Computational Modelling Techniques for Genomics. Nature Reviews Genetics. 20: 389–403.

Fields AP et al. 2015. A Regression-Based Analysis of Ribosome-Profiling Data Reveals a Conserved Complexity to Mammalian Translation. Molecular Cell. 60: 816–827.

Frith MC, Forrest AR, Nourbakhsh E, Pang KC, Kai C, Kawai J, Carninci P, Hayashizaki Y, Bailey TL, and Grimmond SM. 2006. The Abundance of Short Proteins in the Mammalian Proteome. PLOS Genetics. 2: e52.

Goel N, Singh S, and Aseri TC. 2020. Global Sequence Features Based Translation Initiation Site Prediction in Human Genomic Sequences. Heliyon. 6: e04825.

Ji Y, Zhou Z, Liu H, and Davuluri RV. N.d. DNABERT: Pre-Trained Bidirectional Encoder Representations from Transformers Model for DNA-language in Genome. Bioinformatics.

Jorgensen RA and Dorantes-Acosta AE. 2012. Conserved Peptide Upstream Open Reading Frames Are Associated with Regulatory Genes in Angiosperms. Frontiers in Plant Science. 0:

Jumper J et al. 2021. Highly Accurate Protein Structure Prediction with AlphaFold. Nature. 596: 583–589.

Kabir M, Iqbal M, Ahmad S, and Hayat M. 2015. iTIS-PseKNC: Identification of Translation Initiation Site in Human Genes Using Pseudo k-Tuple Nucleotides Composition. Computers in Biology and Medicine. 66: 252–257.

Kalkatawi M, Magana-Mora A, Jankovic B, and Bajic VB. 2019. DeepGSR: An Optimized Deep-Learning Structure for the Recognition of Genomic Signals and Regions. Bioinformatics. 35: 1125–1132.

Makarewich CA et al. 2018. MOXI Is a Mitochondrial Micropeptide That Enhances Fatty Acid β-Oxidation. Cell Reports. 23: 3701–3709.

Matsumoto A, Pasut A, Matsumoto M, Yamashita R, Fung J, Monteleone E, Saghatelian A, Nakayama KI, Clohessy JG, and Pandolfi PP. 2017. mTORC1 and Muscle Regeneration Are Regulated by the LINC00961-Encoded SPAR Polypeptide. Nature. 541: 228–232.

Mudge JM, Ruiz-Orera J, Prensner JR, Brunet MA, Calvet F, et al. 2022. Standardized Annotation of Translated Open Reading Frames. Nature Biotechnology. 40: 994–999.

Mudge JM, Ruiz-Orera J, Prensner JR, Brunet MA, Gonzalez JM, et al. 2021. A Community-Driven Roadmap to Advance Research on Translated Open Reading Frames Detected by Ribo-Seq. bioRxiv. 2021.06.10.447896.

Parikh A, Täckström O, Das D, and Uszkoreit J 2016. A Decomposable Attention Model for Natural Language Inference. In: Proceedings of the 2016 Conference on Empirical Methods in Natural Language Processing. Austin, Texas: Association for Computational Linguistics, pp. 2249–2255.

Pauli A, Valen E, and Schier AF. 2015. Identifying (Non-)Coding RNAs and Small Peptides: Challenges and Opportunities. BioEssays. 37: 103–112.

Ruiz Cuevas MV et al. 2021. Most Non-Canonical Proteins Uniquely Populate the Proteome or Immunopeptidome. Cell Reports. 34: 108815.

Saeys Y, Abeel T, Degroeve S, and Van de Peer Y. 2007. Translation Initiation Site Prediction on a Genomic Scale: Beauty in Simplicity. Bioinformatics. 23: i418–i423.

Stein CS, Jadiya P, Zhang X, McLendon JM, Abouassaly GM, Witmer NH, Anderson EJ, Elrod JW, and Boudreau RL. 2018. Mitoregulin: A lncRNA-Encoded Microprotein That Supports Mitochondrial Supercomplexes and Respiratory Efficiency. Cell Reports. 23: 3710–3720.e8.

Sundararajan M, Taly A, and Yan Q. 2017. Axiomatic Attribution for Deep Networks. arXiv:1703.01365 [cs].

Thibaud-Nissen F, DiCuccio M, Hlavina W, Kimchi A, Kitts PA, Murphy TD, Pruitt KD, and Souvorov A. 2016. P8008 The NCBI Eukaryotic Genome Annotation Pipeline. Journal of Animal Science. 94: 184–184.

Vaswani A, Shazeer N, Parmar N, Uszkoreit J, Jones L, Gomez AN, Kaiser, and Polosukhin I. 2017. Attention Is All You Need. Advances in Neural Information Processing Systems. 30: 5998–6008.

Vitorino R, Guedes S, Amado F, Santos M, and Akimitsu N. 2021. The Role of Micropeptides in Biology. Cellular and Molecular Life Sciences. 78: 3285–3298.

Wang S, Li BZ, Khabsa M, Fang H, and Ma H. 2020. Linformer: Self-Attention with Linear Complexity. arXiv:2006.04768 [cs, stat].

Wei C, Zhang J, and Xiguo Y. 2021. DeepTIS: Improved Translation Initiation Site Prediction in Genomic Sequence via a Two-Stage Deep Learning Model. Digital Signal Processing. 117: 103202.

Wilkie GS, Dickson KS, and Gray NK. 2003. Regulation of mRNA Translation by 5’- and 3’-UTR-Binding Factors. Trends in Biochemical Sciences. 28: 182–188.

Xiong Y, Zeng Z, Chakraborty R, Tan M, Fung G, Li Y, and Singh V. 2021. Nystr\”omformer: A Nystr\”om-Based Algorithm for Approximating Self-Attention. arXiv:2102.03902 [cs].

Yates A et al. 2016. Ensembl 2016. Nucleic Acids Research. 44: D710–D716.

Ye M, Zhang J, Wei M, Liu B, and Dong K. 2020. Emerging Role of Long Noncoding RNA-Encoded Micropeptides in Cancer. Cancer Cell International. 20: 506.

Young SK and Wek RC. 2016. Upstream Open Reading Frames Differentially Regulate Gene-Specific Translation in the Integrated Stress Response. The Journal of Biological Chemistry. 291: 16927–16935.

Zaheer M et al. 2020. Big Bird: Transformers for Longer Sequences. In: Advances in Neural Information Processing Systems. Ed. by H Larochelle, M Ranzato, R Hadsell, MF Balcan, and H Lin. Vol. 33. Curran Associates, Inc., pp. 17283–17297.

Zhang S, Hu H, Jiang T, Zhang L, and Zeng J. 2017. TITER: Predicting Translation Initiation Sites by Deep Learning. Bioinformatics. 33: i234–i242.

Zien A, Rätsch G, Mika S, Schölkopf B, Lengauer T, and Müller KR. 2000. Engineering Support Vector Machine Kernels That Recognize Translation Initiation Sites. Bioinformatics. 16: 799–807.

Zuallaert J, Kim M, Soete A, Saeys Y, and Neve WD. 2018. TISRover: ConvNets Learn Biologically Relevant Features for Effective Translation Initiation Site Prediction. International Journal of Data Mining and Bioinformatics. 20: 267–284.

## References

W. Chen, P.-M. Feng, E.-Z. Deng, H. Lin, and K.-C. Chou. iTIS-PseTNC: A sequence-based predictor for identifying translation initiation site in human genes using pseudo trinucleotide composition. Analytical Biochemistry, 462:76–83, Oct. 2014. ISSN 0003-2697. doi: 10.1016/j.ab.2014.06.022.

K. Choromanski, V. Likhosherstov, D. Dohan, X. Song, A. Gane, T. Sarlos, P. Hawkins, J. Davis, A. Mohiuddin, L. Kaiser, D. Belanger, L. Colwell, and A. Weller. Rethinking Attention with Performers. arXiv:2009.14794 [cs, stat], Mar. 2021.

N. Goel, S. Singh, and T. C. Aseri. Global sequence features based translation initiation site prediction in human genomic sequences. Heliyon, 6(9):e04825, Sept. 2020. ISSN 2405-8440. doi: 10.1016/j.heliyon.2020.e04825.

M. Kabir, M. Iqbal, S. Ahmad, and M. Hayat. iTIS-PseKNC: Identification of Translation Initiation Site in human genes using pseudo k-tuple nucleotides composition. Computers in Biology and Medicine, 66: 252–257, Nov. 2015. ISSN 0010-4825. doi: 10.1016/j.compbiomed.2015.09.010.

M. Kalkatawi, A. Magana-Mora, B. Jankovic, and V. B. Bajic. DeepGSR: An optimized deep-learning structure for the recognition of genomic signals and regions. Bioinformatics, 35(7):1125–1132, Apr. 2019. ISSN 1367-4803. doi: 10.1093/bioinformatics/bty752.

J. M. Mudge, J. Ruiz-Orera, J. R. Prensner, M. A. Brunet, F. Calvet, I. Jungreis, J. M. Gonzalez, M. Magrane, T. F. Martinez, J. F. Schulz, Y. T. Yang, M. M. Albà, J. L. Aspden, P. V. Baranov, A. A. Bazzini, E. Bruford, M. J. Martin, L. Calviello, A.-R. Carvunis, J. Chen, J. P. Couso, E. W. Deutsch, P. Flicek, A. Frankish, M. Gerstein, N. Hubner, N. T. Ingolia, M. Kellis, G. Menschaert, R. L. Moritz, U. Ohler, X. Roucou, A. Saghatelian, J. S. Weissman, and S. van Heesch. Standardized annotation of translated open reading frames. Nature Biotechnology, 40(7):994–999, July 2022. ISSN 1546-1696. doi: 10.1038/s41587-022-01369-0.

Y. Saeys, T. Abeel, S. Degroeve, and Y. Van de Peer. Translation initiation site prediction on a genomic scale: Beauty in simplicity. Bioinformatics, 23(13):i418–i423, July 2007. ISSN 1367-4803. doi: 10.1093/bioinformatics/btm177.

C. Wei, J. Zhang, and Y. Xiguo. DeepTIS: Improved translation initiation site prediction in genomic sequence via a two-stage deep learning model. Digital Signal Processing, 117:103202, Oct. 2021. ISSN 1051-2004. doi: 10.1016/j.dsp.2021.103202.

S. Zhang, H. Hu, T. Jiang, L. Zhang, and J. Zeng. TITER: Predicting translation initiation sites by deep learning. Bioinformatics, 33(14):i234–i242, July 2017. ISSN 1367-4803. doi: 10.1093/bioinformatics/btx247.

A. Zien, G. Rätsch, S. Mika, B. Schölkopf, T. Lengauer, and K.-R. Müller. Engineering support vector machine kernels that recognize translation initiation sites. Bioinformatics, 16(9):799–807, Sept. 2000. ISSN 1367-4803. doi: 10.1093/bioinformatics/16.9.799.

J. Zuallaert, M. Kim, A. Soete, Y. Saeys, and W. D. Neve. TISRover: ConvNets learn biologically relevant features for effective translation initiation site prediction. International Journal of Data Mining and Bioinformatics, 20(3):267–284, Jan. 2018. ISSN 1748-5673. doi: 10.1504/IJDMB.2018.094781.

